# Progressive impairments in executive function in the APP/PS1 model of Alzheimer’s disease as measured by translatable touchscreen testing

**DOI:** 10.1101/742494

**Authors:** A. Shepherd, J.K.H. Lim, V.H.Y. Wong, A.M. Zeleznikow-Johnston, L. Churilov, C.T.O. Nguyen, B.V. Bui, A.J. Hannan, E.L. Burrows

## Abstract

Executive function deficits in Alzheimer’s disease (AD) occur early in disease progression and may be predictive of cognitive decline. However, no preclinical studies have identified deficits in rewarded executive function in the commonly used APP/PS1 mouse model. To address this, we assessed 12-26 month old APP/PS1 mice on rewarded reversal and/or extinction tasks. 16-month-old, but not 13- or 26-month-old, APP/PS1 mice showed an attenuated rate of extinction. Reversal deficits were seen in 22-month-old, but not 13-month-old APP/PS1 animals. We then confirmed that impairments in reversal were unrelated to previously reported visual impairments in both AD mouse models and humans. Age, but not genotype, had a significant effect on markers of retinal health, indicating the deficits seen in APP/PS1 mice were directly related to cognition. This is the first characterisation of rewarded executive function in APP/PS1 mice, and has great potential to facilitate translation from preclinical models to the clinic.

## Introduction

Alzheimer’s disease (AD) is one of the biggest health challenges, with approximately 50 million people currently diagnosed and rates of disease expected to rise in the next 30 years (Alzheimer’s Disease International, 2015). Alzheimer’s disease is characterised by amyloid plaque build-up, tau hyperphosphorylation, neurodegeneration and cognitive decline (Jack Jr et al., 2013). At late stages of disease, the cognitive impairments are so severe that patients are unable to care for themselves, which puts a disproportionate stress on the healthcare system and the family and friends of those affected. Due to this increasing burden of disease, AD has been the subject of intense research for many years. In this time, numerous animal models of AD have been generated, with many potential treatments rescuing cognitive symptoms in a wide variety of models. However, all these treatments have failed to halt progression of the disease in clinical trials. Most preclinical animal studies focus on the assessment of hippocampal deficits as an outcome of treatment efficacy. In the clinic however, other early cognitive changes are seen in AD patients including those in executive function (EF). A greater focus on executive function in preclinical animal models is warranted in order to improve the predictive power of these models to determine how effective treatments will be in the clinic.

Patients with mild cognitive impairment (MCI) and early AD patients show EF deficits on a wide variety of tasks including Tower of London, Stroop Colour-Word Test, Ravens Coloured Progressive Matrices, Ruff Figural Fluency Test, Colour Trails Test Part B, category fluency, backwards digit span, and the Modified Card Sorting Task (Andriuta et al., 2018; Baudic et al., 2006; Huang, Liu, Chang, & Su, 2017; Ramanan et al., 2017). In these tasks, patients show attenuated task acquisition and performance, indicating prefrontal cortex (PFC) dysfunction (Levy-Gigi, Kelemen, Gluck, & Kéri, 2011). More importantly, EF dysfunction, especially on the Trailmaking Part B test (which is part of the Montreal Cognitive Assessment), is predictive of cognitive decline (Mez et al., 2013) and MCI to AD conversion (Albert, Moss, Tanzi, & Jones, 2001; Ewers et al., 2012; Gomar et al., 2011; Huang et al., 2017; Johnson, Storandt, Morris, & Galvin, 2009). As EF deficits occur early in MCI and AD patients and, importantly, have the potential to predict cognitive decline and MCI to AD conversion, these symptoms are of great clinical interest and to date, have not been the focus of testing treatment efficacy in preclinical animal models.

Amyloid-driven mouse models of AD have been shown to have multiple EF deficits early in their progression. First, many models have shown deficits in reversal learning, which assesses a facet of EF called behavioural flexibility (Chudasama, 2011). Briefly, the animal must first learn one association or rule (i.e. the location of platform in a pool or a stimulus-reward pairing), which is then reversed (i.e. the platform is moved to a different location or a different stimulus is now rewarded). Many AD models have shown reversal impairments in spatial maze-based tasks like Morris water maze (Baruch et al., 2015; Hooijmans et al., 2009; Jankowsky et al., 2005; Marchese et al., 2014; Musilli, Nicolia, Borrelli, Scarpa, & Diana, 2013; Papadopoulos, Rosa-Neto, Rochford, & Hamel, 2013; Stover & Brown, 2012), T water maze (Dong et al., 2005; Filali & Lalonde, 2009; Filali, Lalonde, & Rivest, 2011; Filali et al., 2012), cheeseboard (Cheng, Logge, Low, Garner, & Karl, 2013) and Barnes maze (Stover, Campbell, Van Winssen, & Brown, 2015) as well as olfactory (Girard et al., 2013, 2014; Guérin, Sacquet, Mandairon, Jourdan, & Didier, 2009), digging (Shirey et al., 2009; Zhuo et al., 2008, 2007) and place preference (Masuda et al., 2016) reversal tasks. AD models have also shown deficits in instrumental extinction, another test of EF, where the animal must learn to stop responding to a conditioned stimulus (i.e. a stimuli that previously indicated a reward or punishment). Extinction tasks are similar to reversal tasks in that an animal must learn to adjust a stimulus/outcome association, but without having to learn a new rule. AD models also show enhanced extinction in rewarded tasks (Romberg, Horner, Bussey, & Saksida, 2013) and a mix of attenuated and enhanced extinction in aversive tasks like fear conditioning (Bonardi, de Pulford, Jennings, & Pardon, 2011; Cheng et al., 2019) and conditioned taste aversion (Hanna et al., 2009; Janus et al., 2004; Pardon et al., 2009; Ramírez-Lugo, Jensen, Søderman, & West, 2009; Rattray et al., 2010; Rattray, Scullion, Soulby, Kendall, & Pardon, 2009). One tau model has also been shown to have enhanced extinction on conditioned taste aversion (Pennanen, Welzl, D’Adamo, Nitsch, & Götz, 2004). In APP/PS1 mice, fear extinction was impaired while appetitive extinction was left intact, indicating that aversive extinction is impaired earlier than rewarded extinction (Bonardi et al., 2011). However, the vast majority of tasks applied to AD mouse models either rely on aversive environments or ethologically relevant mouse behaviour; thus they are not analogous to those performed clinically, where the patient is voluntary responding to stimuli on paper or a screen. The dissociation between rewarded and aversive extinction highlights the need for preclinical models to mirror the clinic as closely as possible, to improve testing and translation of treatments. As such, we need a paradigm that can assess EF in non-aversive, human relevant conditions.

Touchscreen testing is a relatively new paradigm to behaviourally test AD mouse models with great translational potential (Shepherd, Tyebji, Hannan, & Burrows, 2016). To date, two AD mouse models, the TgCRND8 and APPSwDI/Nos2-/- mice, have undergone characterisation of EF using reversal and extinction paradigms in touchscreens. Contrary to predictions from previously used aversive tasks, TgCRND8 mice show both accelerated reversal and extinction in touchscreens compared to WT mice. This is not due to an accelerated ‘forgetting’ of the first stimulus/reward association, as TgCRND8 mice show intact retention memory for the initial stimulus (Romberg et al., 2013). Conversely, APPSwDI/*Nos2-/-* mice showed an attenuation of reversal in the touchscreen testing paradigm (Piiponniemi et al., 2017). One of the most commonly characterised AD mouse models, the APP/PS1 mouse has not yet been characterised on touchscreen-based EF tasks. Given previous reports showing slower acquisition of reversal Morris water maze (MWM) (Hooijmans et al., 2009; Jankowsky et al., 2005; Stover & Brown, 2012) and left-right discrimination learning (Filali, Lalonde, & Rivest, 2009; Filali et al., 2011) in APP/PS1 mice, we aimed to assess whether reversal learning was impaired in this mouse model using the pairwise discrimination and reversal task. Conflicting findings exist for extinction learning in APP/PS1 mice, with reports of both attenuated (Bonardi et al., 2011) and enhanced extinction to aversive tasks (Cheng et al., 2019; Ramírez-Lugo et al., 2009) in the mouse model. When assessed in positively rewarded extinction tasks, APP/PS1 mice did not demonstrate any deficits (Bonardi et al., 2011; Cheng et al., 2013). To resolve this conflict, we set out to test if APP/PS1 mice would show deficits in reward-based EF tasks using the rodent touchscreen paradigm. We hypothesised that APP/PS1 mice would develop EF deficits in both appetitive reversal and extinction tasks.

## Methods

### Animals

APP/PS1 mice (APPswe/PS1ΔE9 on a B6C3 hybrid background, Jax strain #4462) were obtained from Jackson Laboratory and bred onsite in the SPF facility at the Florey Institute of Neuroscience and Mental Health (Parkville, VIC, Australia). Male APP/PS1 and wild type littermates (WT) mice from 12-26 months of age were used in experimental procedures. In the results sections, groups of animals are referred to by the rounded average age of the group. Females were not tested due to interference with male performance and lack of single-sex equipment. Animals were housed in individually ventilated cages with *ad libitum* food and water until experimental procedures began. Prior to the start of experimental procedures, animals were moved to food restricted, open top housing with a reversed light cycle (12:12 hour, 7am lights off), at 20°C ± 1°C. Animals were housed in groups of 2-4 unless fighting necessitated single housing, and all cages had a small shelter and bedding material. Cohort 1 and cohort 3 animals were initially restricted to 85% free feeding weight (FFW), while cohort 2 was restricted to 75% FFW. A stricter regime was applied to cohort 2 as they started food restriction at 18 months of age, where the average weight was over 40 g, and 85% FFW was insufficient to motivate animals to complete the total number of trials each day. All procedures were approved by The Florey Institute of Neuroscience and Mental Health Animal Ethics Committee (AEC # 16-020) and were conducted in accordance with the Australian Code of Practice for the Care and Use of Animals for Scientific Purposes as described by the National Health and Medical Research Council of Australia.

### Apparatus

Behavioural testing was undertaken in automated touchscreen-based operant systems (Campden Instruments Ltd, Loughborough, UK). Automated software Whisker Server and ABET II were used to run the tasks and collect the data respectively (Lafayette Instruments, Lafayette, IN, USA). Experimenters were blinded to genotype throughout experimental procedures, and the apparatus cleaned (including the screen) between each animal with 70% ethanol.

### Behavioural Procedures

#### Pre-training

Pre-training procedures have been described in detail previously (Mar et al., 2013). All behavioural testing was conducted under red light during the dark phase of the animals. Animals were initially habituated to a reward (Iced Strawberry Milk, Nippy’s Ltd, Moorook, SA, Australia) in their home cage for 2 days, then habituated to the touchscreen chambers with freely available reward for 2 days. Mice were then trained to nose-poke stimuli on a screen, for which they were rewarded 7 µL per trial. Following this, animals were required to learn to initiate 30 trials in an hour by placing their head into the food magazine. Finally, animals were trained to touch only the stimulus by having a light turned on and a 5s time out initiated if they touched the blank parts of the screen. The inter-trial interval was 5s during all stages. During this final stage, animals were required to touch the stimulus rather than the blank parts of the screen for 80% of the 30 trials.

#### Pairwise Discrimination and Reversal

Pairwise discrimination (PD) and reversal task training have been described in detail previously (Mar et al., 2013). In the PD task, animals were required to self-initiate trials by placing their head in the food magazine. Following this, animals were presented with two opposite diagonal contrast-grating images (45° and 120°); one of which was rewarded (correct) and one of which was not (incorrect). The location of the correct image was pseudorandomised between trials, ensuring the rewarded image was not presented in the same location more than 3 times consecutively. A nose-poke to the correct stimulus caused a tone to sound and delivery of a 7µL reward. Conversely, choosing the incorrect stimulus caused both a house light and a 5s timeout to trigger, analogous to the final stage of pre-training. However, in this case, a correction trial loop was started; during correction trials, the same trial was presented repeatedly until the animal fixed their initial mistake. Animals were required to perform 30 unique (non-correction) trials in one hour, and animals were considered to have learnt the discrimination if they chose the correct image for 80% of unique trials for 2 consecutive days. Once animals reached criterion, they were rested (with weekly reminder sessions) until all animals in the cohort had finished PD. Following PD, animals all started reversal; the procedure was the same as PD, except the reward contingencies for the images are reversed i.e. the previously incorrect image was rewarded while the previously correct image was punished. Animals were required to inhibit their response to the previously rewarded image and learn a new stimulus/reward association to the previously incorrect image. Criterion for reversal was 75% correct for unique trials for 2 consecutive days. Correct stimuli were pseudo-randomly assigned prior to experimental initiation, ensuring equal balance across genotypes.

#### Contrast Probe

The contrast probe has not been previously described. Following reversal, animals were moved to the contrast probe, which had both an acquisition and probe stage. In the acquisition phase, animals were presented with two images; one unrewarded grey square and one rewarded contrast-grating. The rewarded image was kept consistent with the correct image assigned to the animal in reversal. The animal was required to finish 30 trials at 80% correct to move to the probe phase. In the probe, animals initiated a trial and were presented with 2 images; one unrewarded grey square, and a contrast-grating image at one of four levels of contrast (although with matching levels of luminance; 100%, 50%, 25% and 12.5% (Kim et al., 2015). Level of contrast used in each trial was pseudorandomly determined, ensuring equal presentation of all 4 contrast images per session. Animals were rewarded for correctly choosing the contrast-grating image compared to the grey square. Animals were run for 4 days on the contrast probe.

#### Extinction

Extinction in rodent touchscreens has been previously described (Mar et al., 2013), and was always performed as the last task the animals undertook. Briefly, the extinction task had two distinct phases; acquisition and extinction. In acquisition, the animal had to associate touching a white square stimulus and tone with an immediate reward. Animals were required to initiate and complete 30 trials in 13 minutes or less for 5 consecutive days. In the extinction phase, the reward is removed, so that when animals touched the square, the reward tone was still played but no reward was delivered. The square was automatically presented for 30 seconds per trial, 30 trials a day, for seven days.

### Study design

The behavioural components of this study were conducted over 3 cohorts of APP/PS1 animals at various ages, which are summarised in Table 1. For all experiments, researchers were blind to genotype, with behavioural data automatically collected and analysed using previously published methods, thus eliminating bias. Sample sizes of approximately 10-12 per group were used across experiments. APP/PS1 animals showed a higher premature death rate than their WT littermate counterparts, leading to a smaller number in this group both at the start and end of experiments.

**Table 1.** Numbers and ages of wildtype (WT) and APPswe/PS1ΔE9 (APP/PS1) animals used in each behavioural task.

| Animal Cohort | Task | Genotype |  | Age (months) | Age started food restriction |
| --- | --- | --- | --- | --- | --- |
|  |  | WT | APP/PS1 |  |  |
| Cohort 1 | PD | 10 | 9 | 12-12.5 | 6 months |
|  | Reversal | 10 | 9 | 12.5-13 |  |
|  | Extinction | 10 | 8 | 13-13.5 |  |
| Cohort 2 | PD | 12 | 9 | 21-22 | 18 months |
|  | Reversal | 12 | 8 | 22-25.5 |  |
|  | Contrast probe | 12 | 6 | 22-25.5 |  |
|  | Extinction | 10 | 7 | 25.5-26.5 |  |
| Cohort 3 | Extinction | 12 | 8 | 16-16.5 | 8 months |

This study included 3 cohorts of animals, ranging in age from 12 to 26.5 months of age. Animal numbers for up to 4 behavioural tasks are broken down by genotype and age for each distinct cohort of animals. WT = Wildtype, PD = Pairwise Discrimination

#### Visual testing

##### Electroretinography (ERG)

ERG procedures have been described in detail for mice previously (Zhao et al., 2017). Three groups of animals underwent ERG; 12-month-old touchscreen trained APP/PS1 and WT mice (cohort 1; APP/PS1, n = 9; WT, n = 11), 20-26 month old mice (cohort 2; APP/PS1, n = 7; WT, n = 9), and a separate cohort of 20-26 month old behaviourally naïve, standard housed mice (cohort 3; APP/PS1, n = 6; WT, n = 8). This latter group was used to assess if touchscreen training altered visual function. Briefly, mice were dark-adapted in a well-ventilated light-proofed room for at least 12 hours prior to ERG recording to maximise light sensitivity. Immediately before measurements, animals were anesthetised with ketamine/xylazine in saline (80 mg/kg and 100 mg/kg, respectively from Ilium, Troy Laboratory Pty Ltd, Smithfield, NSW, Australia) and, once unconscious, were administered with one drop of topical anaesthesia (proxymetacaine, 0.5% Alcaine^TM^, Alcon Laboratories, Frenchs Forest, NSW, Australia) and one drop of a mydriatic tropicamide 1% (Mydriacyl^TM^, Alcon Laboratories) to each eye. The animal was then strapped to a water-heated platform to prevent anaesthesia induced temperature loss and a drop of ocular lubricant (Celluvisc, carmellose sodium 10 mg/mL, Allergan, Gordon, NSW, Australia) was applied. Active and inactive electroplated (chlorided) silver electrodes were placed on the corneal surface of the eye (active) and around the eye behind the limbus (reference), while a ground electrode was inserted into the tail. All ERG preparation procedures were performed in darkness with the aid of dim red headlamps. Once prepared, animals were placed directly under a Ganzfeld sphere (Photometric Solutions International, Huntingdale, VIC, Australia) within a custom-built Faraday cage. After a brief re-adaptation period of 10 minutes, recordings were made in response to brief flashes of light starting with dim and progressing to brighter intensities (from −6.35 to 2.07 log cd.s/m^2^). Analysis of waveforms was performed as described previously (Nguyen et al., 2016).

##### Optical coherence tomography (OCT)

Following ERG measurements, animals were placed on a heat pad for 10 minutes with Genteal gel and small glass coverslips placed on each eye to clear anaesthesia-induced cataracts. At this stage, animals were administered a 25% dose anaesthetic top up of ketamine/xylazine as described above, if required. OCT scans were acquired using a spectral domain OCT (Spectralis OCT, Heidelberg Engineering, Heidelberg, Germany) with a 25D imaging lens attached (adapted for mouse retinal imaging), with the mouse placed on a rigid animal platform (Thorlabs Inc. Newton, NJ, USA).. A volume scan (7.6 × 6.3 × 1.9 mm) centred on the optic nerve head consisting of 121 evenly spaced horizontal B-scans, each consisting of 768 A-scans. Axial resolution and lateral resolution was 3.87 µm and 9.8 µm per pixel, respectively. Throughout imaging, corneal hydration was maintained using lubricating eyedrops (Systane^TM^ Ultra, Alcon Laboratories). B-scans were exported as TIFFs and were analysed by a masked researcher as described previously (Zhao et al., 2017). Retinal nerve fibre layer (RNFL) was measured from the posterior vitreous interface to the anterior inner plexifrom layer while total retinal thickness (TRT) was measured from the vitreous interface to Bruch’s membrane.

### Statistical methods

The effect of genotype on number of trials to criterion and number of correction trials to criterion in PD and reversal were analysed at the animal level using a Log-rank Mantel Cox test and unpaired Student’s t-test with Welsh’s correction, respectively. To investigate percent responses in extinction, two-way repeated measures ANOVA’s were utilised, with genotype and day as factors. To further investigate the effect of genotype and day in touchscreen tasks, we used generalised linear, latent and mixed models (GLLAMM, package from (Rabe-Hesketh, Skrondal, & Pickles, 2005)) of regression on a trial-by-trial level, using individual animals as random effects to reflect clustered nature of all observations within each animal, as previously described (Zeleznikow-Johnston, Burrows, Renoir, & Hannan, 2017). For investigating odds of correct responses (for PD, reversal and the contrast probe) or responses overall (extinction), random effect logistic regressions were used with corresponding effect sizes reported as adjusted odds ratios (aORs) with respective 95% confidence intervals (95% CIs). For interpretation, an aOR of smaller than 1 indicated that APP/PS1 mice were less likely to have responded correctly to a trial, while the corresponding 95% CI not including 1 represented statistical significance (p<0.05). When investigating correction trial counts for PD or reversal, random effect Poisson regressions were used with corresponding effect sizes reported as adjusted incidence rate ratios (aIRRs) with respective 95% CIs. If the aIRR was under 1, this indicated that the APP/PS1 mice had a lower expected count of correction trials compared to WT animals. As for the aOR, the corresponding 95% CI not including 1 represented statistical significance (p < 0.05).

For analysis of retinal nerve fibre layer thickness, total retinal thickness and amplitude of response of photoreceptor, bipolar cells, retinal ganglion cells and amacrine cells, one-way ANOVA’s with Welch correction was used. Here, each data point represented the average of the two eyes of each animal and the group sizes were as follows: Naive WT: n = 8, naive APP/PS1: n = 7, 24 month touchscreen trained (TS) animals from cohort 2, WT: n = 9 and APP/PS1: n = 6, and 12 month old TS animals, WT: n = 11 and APP/PS1: n = 9. All GLLAMM statistical analyses were conducted using STATA v13IC (StataCorp, College Station, TX USA), with all other analyses conducted with SPSS v26 (IBM, Chicago, IL USA).

## Results

### Pairwise discrimination is unaffected in 12- and 21-month-old APP/PS1 mice

For PD, animals were trained to choose one diagonal contrast-grating image over another for a strawberry milk reward. At 12 months of age, APP/PS1 animals took the same number of trials to learn the task (Figure 1A, χ2 (1) = 0.05, p = 0.82) and, when analysed at the trial level, were no less likely to choose the rewarded image than their WT counterparts given any unique trial (Figure 1C, aOR = 0.97, (95% CI = 0.67; 1.39), p = 0.85). Similarly, 21-month-old APP/PS1 mice took approximately the same number of trials to learn the discrimination (Figure 1B, χ2(1) = 2.71, p = 0.10), and, when analysed at the trial level, the odds of correct selection were unaffected by genotype (Figure 1D, aOR = 1.04, (95% CI = 0.70; 1.55), p = 0.83). If animals chose the incorrect image, they were exposed to the same image and location, termed a correction trial. Correction trials are indicative of perseveration. Overall, there was no difference in the number of correction trials to criterion in 12-month-old (Figure 1E, t(12.76) = 0.65, p = 0.53) or 21-month-old (Figure 1F, t(9.95) = 1.50, p = 0.16) APP/PS1 and WT mice. Similarly, when analysed at the level of trial, the expected number of correction trials (per incorrect trial), as estimated by the incidence rate ratio, was unaffected by genotype in both 12-month-old (Figure 1G, aIRR = 1.031, (95% CI = 0.86; 1.23), p = 0.74) and 21-month-old animals (Figure 1H, aIRR = 0.96, (95% CI = 0.67; 1.37), p = 0.81).

**Figure 1.**
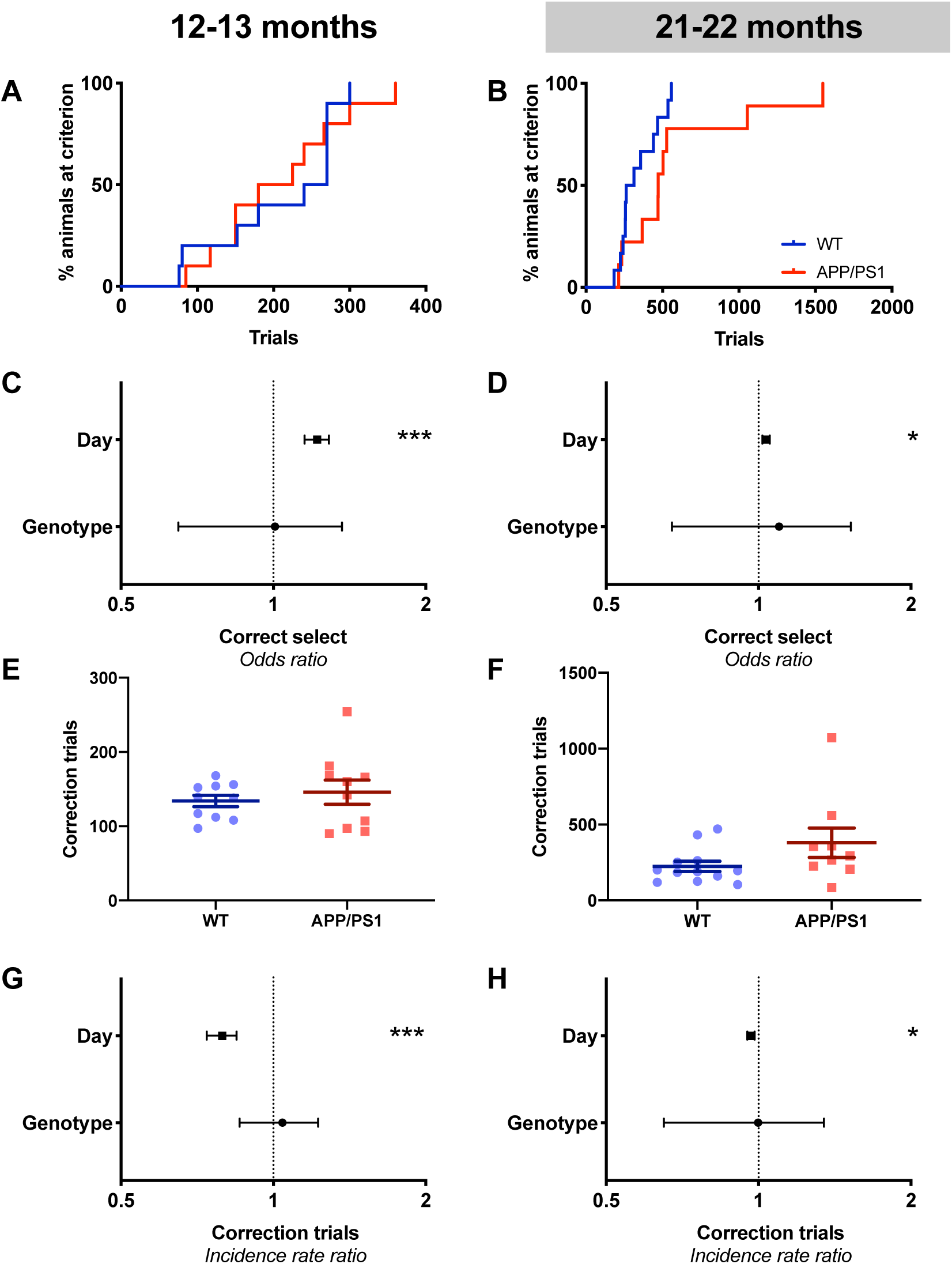
Pairwise discrimination was unaffected in 12- and 21-month-old APP/PS1 mice. 12-month-old (A) and 21-month-old (B) old APP/PS1 mice took an equivalent number of trials to reach criterion. Similarly, the odds of correct selection were not affected by genotype in 12- (C) or 21- (D) month-old animals, although in both groups day significantly increased the odds of correct selection. 12- and 21- month-old APP/PS1 mice completed an equivalent number of correction trials as their WT counterparts, both in overall number (E, F), and in the rate of correction trials they do per incorrect trial (G, H). Day significantly decreased the incidence rate ratio of correction trials in both age groups (G, H). A-B show survival curves, E-F show mean ± SEM while C-D and G-H show odds ratios ± 95% CI. * = p < 0.05; *** = p < 0.001

Behavioural performance improved over days, as shown by the increased odds of correct selection (Figure 1C, 12-month-old aOR = 1.22 (95% CI = 1.15; 1.29), p<0.001, Figure 1D, 21-month-old aOR = 1.04 (95% CI = 1.00; 1.05), p<0.001) and decreased expected count of correction trials (Figure 1G, 12-month-old aIRR = 0.790, (95% CI = 0.74; 0.85), p < 0.001, Figure 1H, 21-month-old aIRR = 0.97, (95% CI = 0.95; 0.98), p<0.001), indicating intact learning in all groups. Age also had a significant effect on task measures (Supplementary Figure 1), with 21-month-old WT animals taking more trials to reach criterion (Supplementary Figure 1A, χ2(1)=4.24, p=0.038), performing more correction trials overall (Supplementary Figure 1B, t(12.11) = 2.61, p = 0.023) and showing a decreased odds of correctly selecting the rewarded image given any unique trial (Supplementary Figure 1C, aOR= 0.53, (95% CI = 0.37; 0.76), p = 0.001) when compared to 12-month-old WT mice. The expected count of correction trials per incorrect trial remained unchanged between the two WT age groups (Supplementary Figure 1D, aIRR= 1.08, (95% CI= 0.76; 1.53), p=0.69).

### 22-month-old, but not 13-month-old APP/PS1 mice show impairments in reversal learning

Following PD, the reward contingencies were swapped, and reversal learning to criterion was assessed. 13-month-old APP/PS1 mice took a similar number of trials as their WT counterparts to reach criterion (Figure 2A, χ2(1)= 0.21, p=0.64), and were just as likely to choose the correct, rewarded image given any unique trial (Figure 2C, aOR = 0.98, (95% CI = 0.69; 1.40), p = 0.93). In comparison, 22-month-old APP/PS1 mice took significantly more trials to reach criterion (Figure 2B, χ2(1) = 9.70, p < 0.001). This impairment was also reflected at the level of trial, with 22-month-old APP/PS1 mice showing reduced odds of correct selection (Figure 2D, aOR = 0.72, (95% CI = 0.54; 0.96), p = 0.025). 13-month-old APP/PS1 animals perform a similar number of correction trials to criterion (Figure 2E, t(13.55) = 1.15, p = 0.27) and an equivalent expected count of correction trials per incorrect trial (Figure 2G, aIRR = 0.98, (95% CI = 0.69; 1.40), p = 0.93) to their WT counterparts. As would be expected, 22-month-old APP/PS1 mice showed an increased number of correction trials overall (Figure 2F, t(9.94) = 3.15, p = 0.0052) as well as an increased expected count per incorrect trial (Figure 2H, aIRR = 1.35, (95% CI = 1.23; 1.48), p < 0 .001); the latter indicating APP/PS1 mice take longer to correct mistakes compared to their wild type counterparts. All animals reached criterion. To ensure this phenomenon was not due to the APP/PS1 mice completing less trials than their WT counterpart, which would in turn reduce the opportunities to learn the task. There was no effect of genotype on the number of unique trials (Supplementary Figure 2A, F(18,1) = 1.959, p=0.52), correction trials (Supplementary Figure 2B, F(18,1) = 1.714, p=0.55) or the combined number of trials (Supplementary Figure 2C, F(18,1) = 11.46, p=0.23). Thus, the deficit seen here is driven by a cognitive, not motivational, phenotype in the APP/PS1 mice.

**Figure 2.**
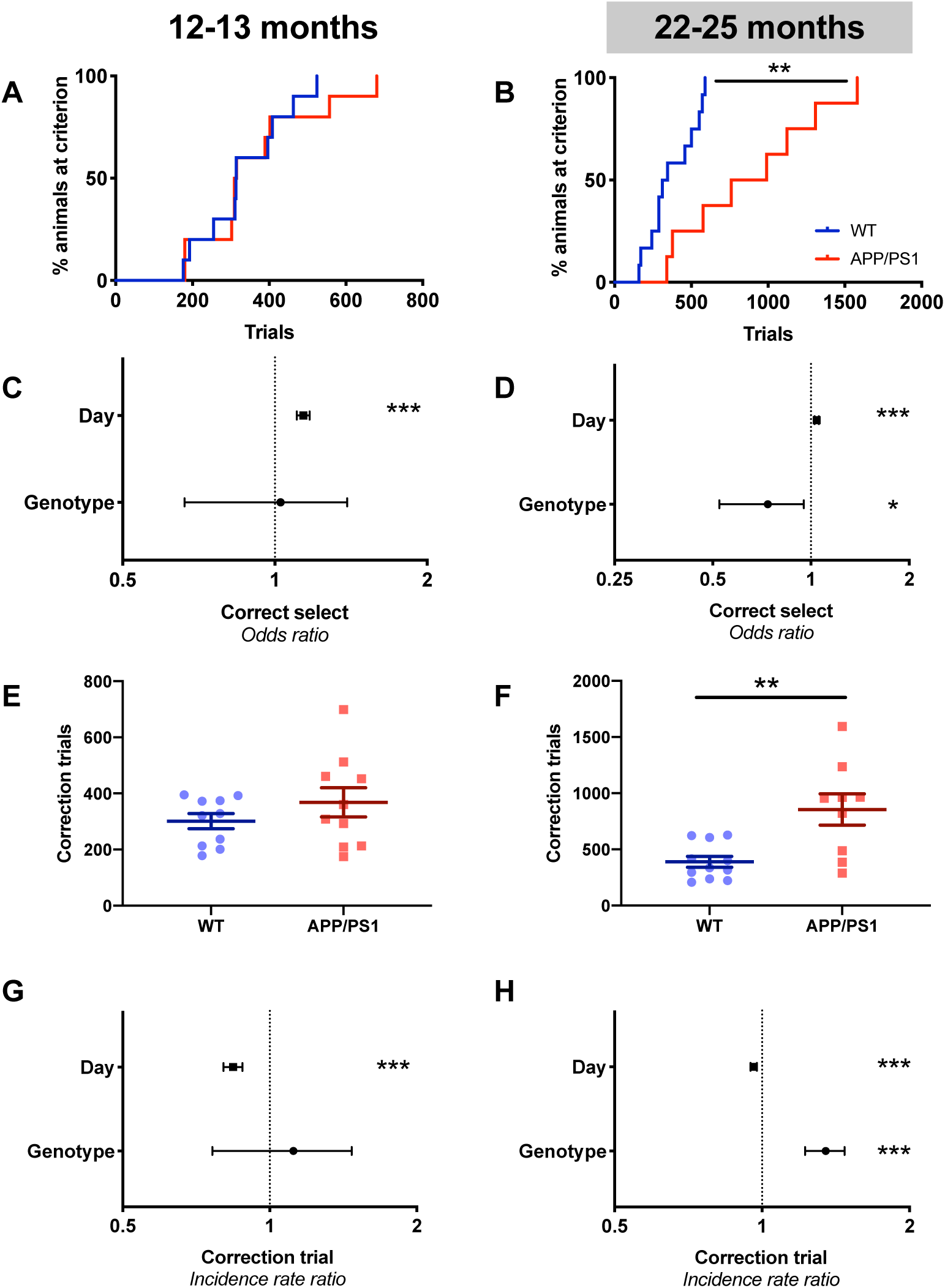
22-month-old APP/PS1 mice were impaired in reversal learning, while 13-month-old mice were unaffected. 13-month-old APP/PS1 mice took an equivalent number of trials to reach criterion for reversal and were just as likely to choose the correct image given any unique trial (A, C). Conversely, 22-month-old APP/PS1 mice took longer to reach criterion (B) and were less likely to choose the correct image given any unique trial (D). Day increased the odds of correct selection at both ages (C-D). 13-month-old APP/PS1 mice completed an equivalent number of correction trials overall as their WT counterparts (E) and showed the same expected correction trial count per incorrect trial as WT animals (G). 22-month-old APP/PS1 mice performed significantly more correction trials than their WT counterparts (F) and performed more correction trials per incorrect trial than WT animals (H). Day decreased the incidence rate ratio of correction trials at both ages (G-H). A-B show survival curves, E-F show mean ± SEM while C-D and G-H show odds ratios ± 95% CI. * = p < 0.05; *** = p < 0.001

Reversal learning can also be split into two stages; initially, animals had to learn to inhibit their responding to the original reward-stimulus association (early reversal, where the percent correct is below chance (50% per session)) and then learn the new reward-stimulus association (late reversal; where percent correct is above chance). When assessing early and late reversal, similar trends were seen to overall reversal, regardless of the stage. 13-month-old APP/PS1 and WT animals showed equivalent odds of correct selection at early (Supplementary Figure 3A, aOR = 0.94, (95% CI = 0.79; 1.12), p = 0.49) and late reversal (Supplementary Figure 3E, aOR = 0.94, (95% CI = 0.73; 1.20), p = 0.60) while 22-month-old APP/PS1 animals showed a non-significant decrease in the odds of correct selection at early (Supplementary Figure 3B, aOR = 0.79, (95% CI = 0.61; 1.03), p = 0.078) and late (Supplementary Figure 3F, aOR = 0.82, (95% CI = 0.65; 1.03), p = 0.082) reversal. In terms of correction trials, 13 months old APP/PS1 mice showed similar expected counts of correction trials at both early (Supplementary Figure 3C, aIRR = 1.11, (95% CI = 0.87; 1.43), p = 0.388) and late reversal (Supplementary Figure 3G, aIRR= 1.15, (95% CI= 0.86; 1.54), p = 0.35). In 22-month-old APP/PS1 mice, the increased rate of correction trials was significant only for early (Supplementary Figure 3D, aIRR = 1.21, (95% CI = 1.01; 1.44), p = 0.04) not late (Supplementary Figure 3H, aIRR = 1.31, (95% CI = 0.97; 1.78), p = 0.081) reversal, indicating the APP/PS1 animals were impaired during early reversal.

Despite impairments in the older mice, learning over time during the reversal task was evident in both 13-month-old and 22-month-old mice. This was evident in the increase odds of correct selection over days (Figure 2C, 13-month-old aOR = 1.14, (95% CI = 1.11; 1.17), p<0.001, Figure 2D, 22-month-old aOR = 1.042, (95% CI = 1.027; 1.058), p < 0.001) and decreasing the expected count of correction trials (Figure 2G, 13-month-old aIRR = 0.84, (95% CI = 0.80; 0.88), p < 0.001, Figure 2H, 22-month-old aIRR = 0.96, (95% CI = 0.95; 0.98), p < 0.001). Age also had a significant effect at the trial level, with 22-month-old WT animals showing decreased odds of correctly selecting the rewarded image given any unique trial (Supplementary Figure 1G, aOR= 0.56, (95% CI = 0.43; 0.73), p < 0.001) and an increased expected count of correction trials per incorrect trial (Supplementary Figure 1H, aOR = 1.72, (95% CI = 1.39; 2.12), p < 0.001) compared to 13-month-old WT animals. Conversely, the number of trials (Supplementary Figure 1E, χ2(1) = 0.99, p = 0.32) and correction trials (Supplementary Figure 1F, t(15.49) = 1.59, p = 0.13) to criterion were equivalent in the two age groups of WT mice.

### 16-month-old APP/PS1 mice show extinction impairments

Extinction tested an animal’s reduction in response to a stimulus when presented without reward. Extinction was performed in 3 cohorts of animals at 13, 16 and 26 months of age. All animals completed the acquisition stage prior to extinction, excepting two 26-month-old animals. Regardless of age and genotype, all animals showed a decreased odds of response over the 7 days of extinction, indicating that the animals learnt that touching the stimulus no longer caused reward delivery (Figure 3D, 13 months: aOR= 0.64, (95% CI= 0.59; 0.69), p<0.001, Figure 3E, 16 months: aOR= 0.63, (95% CI= 0.57; 0.69), p<0.001, Figure 3F, 26 months: aOR= 0.804, (95% CI= 0.77; 0.84), p<0.001). At 13 months of age, a difference in mean total responses over each day was seen between APP/PS1 and WT mice over the 7 days of extinction (Figure 3A, F(6,17)=15.62, p=0.001). Similarly, 16-month-old APP/PS1 showed a higher mean total response over the 7 days of extinction (Figure 3B, F(6, 18)=8.65, p=0.0087). Older 26-month-old APP/PS1 mice did not show any difference in mean total response to the stimulus over time (Figure 3C, F(6,11)=0.11, p=0.75). We then analysed these responses at the trial level, and while 1-month-old APP/PS1 mice were significantly more likely to respond (Figure 3D, aOR= 1.90, (95% CI= 1.35; 2.67), p<0.001), this remained constant over time, reflected by a non-significant genotype-day interaction (Figure 3D, aOR= 0.99, (95% CI= 0.85; 1.15), p=0.90). This indicated that the rate of extinction was equivalent between APP/PS1 and WT mice. At 16 months of age, APP/PS1 mice showed increased odds of responding (Figure 3E, aOR= 1.75, (95% CI= 1.25; 2.46), p<0.001) and compared to WT mice, this did not decline as quickly over time, as evidenced by a significant genotype-day interaction (Figure 3E, aOR= 1.20, (95% CI= 1.02; 1.41), p=0.025). This indicated a difference in the rate of extinction between APP/PS1 and WT mice at this age. This effect was lost by 26 months of age, with no genotype (Figure 3F, aOR= 1.04, (95% CI= 0.64; 1.70), p=0.87) or a genotype-day interaction effect (Figure 3F, aOR= 1.04, (95% CI= 0.94; 1.14), p=0.478). The interpretation of extinction at 26 months of age is confounded by the fact that mice are significantly slower to respond to the white square compared to 13- and 16-month-old animals (data not shown: median difference (s)= −0.64 (95% CI= −1.17; −0.10), p=0.02). Both 26-month-old mice groups showed significantly lower responses (responding to approximately 65.05% of trials) on day 1 compared to 13 and 16-month-old animals (responding to approximately 85.25% and 79.45% of trials respectively) (Figure 3A-C, F(2,43)=6.92, p=0.0022). This lack of engagement with the task from the beginning of extinction learning may have introduced a floor effect, masking any genotype differences.

**Figure 3.**
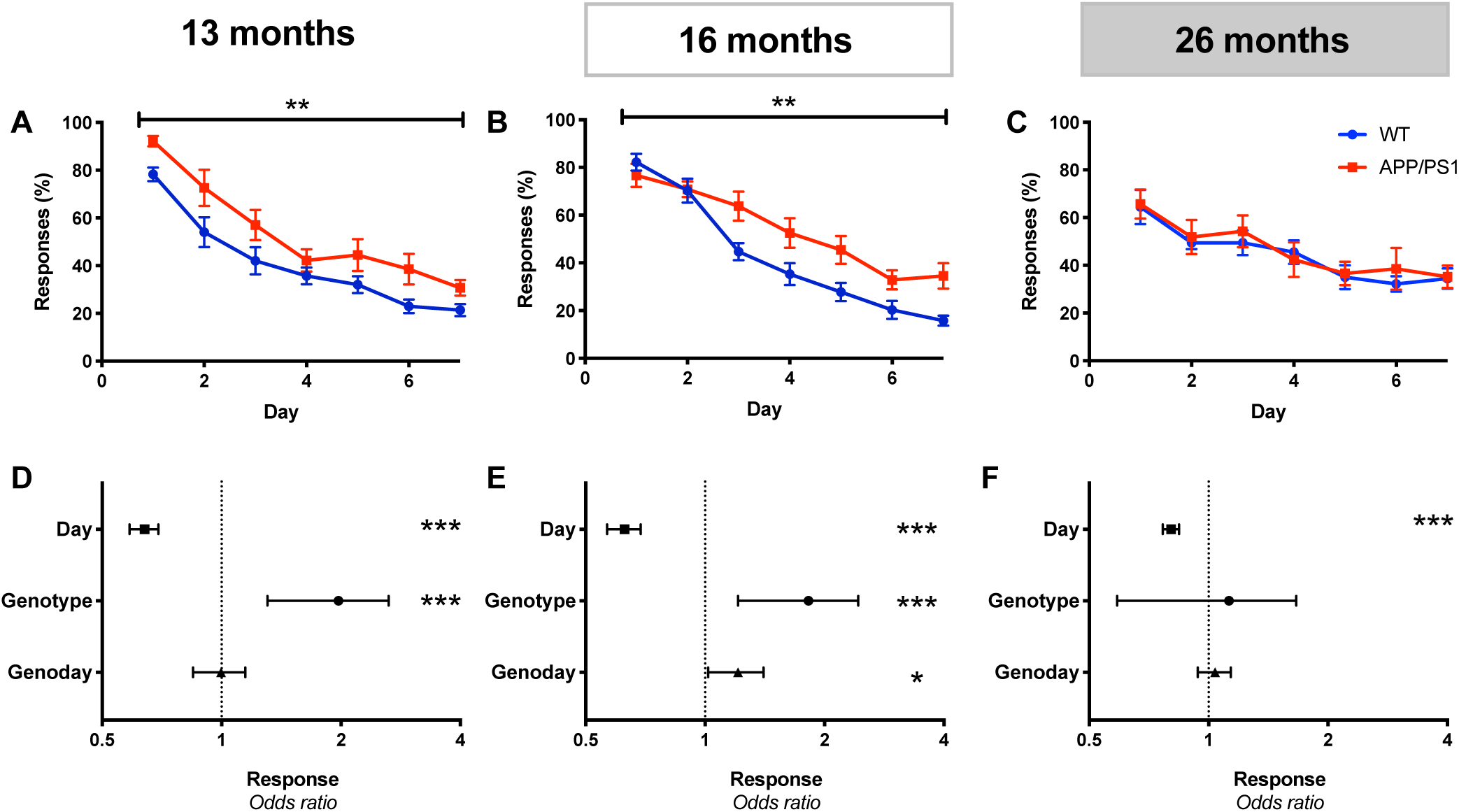
16-month-old APP/PS1 mice showed extinction deficits. 13- and 16-month APP/PS1 mice showed genotype effects over extinction, with only 16-month-old mice showing an attenuated rate of response. 26-month-old mice showed no genotype effects over extinction. 13-month-old APP/PS1 mice showed an increased response from day 1 (A) while 16-month-old APP/PS1 mice had an increased response at day 3 (B). 26-month-old mice showed an equivalent change in responses over days between genotypes (C). 13-month-old APP/PS1 mice had significantly increased odds of response but no effect of a genotype day interaction variable, indicating the rate of extinction was equivalent (D). 16-month-old APP/PS1 mice showed increased odds of response and a significant genotype day interaction effect, indicating that the rate of extinction was different between APP/PS1 and WT mice (E). 26-month-old APP/PS1 mice showed neither a genotype nor a genotype day interaction effect on the odds of making response during extinction (F). All ages showed decreased odds of response over days, indicating inhibition of the old stimulus/reward contingency (G-I). A-F show mean ± SEM and G-I show odds ratios ± 95% CI. * = p < 0.05; ** = p < 0.01; *** = p < 0.001

### 26-month-old mice show impaired retinal reactivity regardless of genotype

Previous reports have indicated that APP/PS1 mice at 20-26 months of age have impaired vision which could account for differences in touchscreen performance (Stover & Brown, 2012). As such, we wished to test the vision of touchscreen trained 12-month-old and 22-26 month old APP/PS1 and WT animals, in addition to comparing touchscreen trained animals to a cohort of behaviourally naïve 20-26 month old animals.

First, following the reversal task, the 23-month-old APP/PS1 animals were subjected to a modified two-choice discrimination task to test vision in the touchscreens apparatus. In this task animals were shown a grey square or a contrast grating of one of four contrasts – 100%, 50%, 25% or 12.5%, and rewarded for selecting the contrast-grating. As contrast was decreased, the odds of an animal correctly selecting the rewarded contrast-grating decreased (Figure 4A-B, aOR = 9.21, (95% CI = 5.13; 16.53), p < 0.001). Performance on the task was unaffected by genotype (Figure 4A-B, aOR = 0.95, (95% CI = 0.55; 1.65), p = 0.86), indicating that APP/PS1 animals could perceive all contrast images just as well as their WT counterparts. While it is unlikely the reduced performance of the APP/PS1 in the reversal task was attributable to visual changes, we further subjected animals to optical coherence tomography (OCT) and electroretinography (ERG) to assess structural and functional integrity of the retina to confirm this.

**Figure 4.**
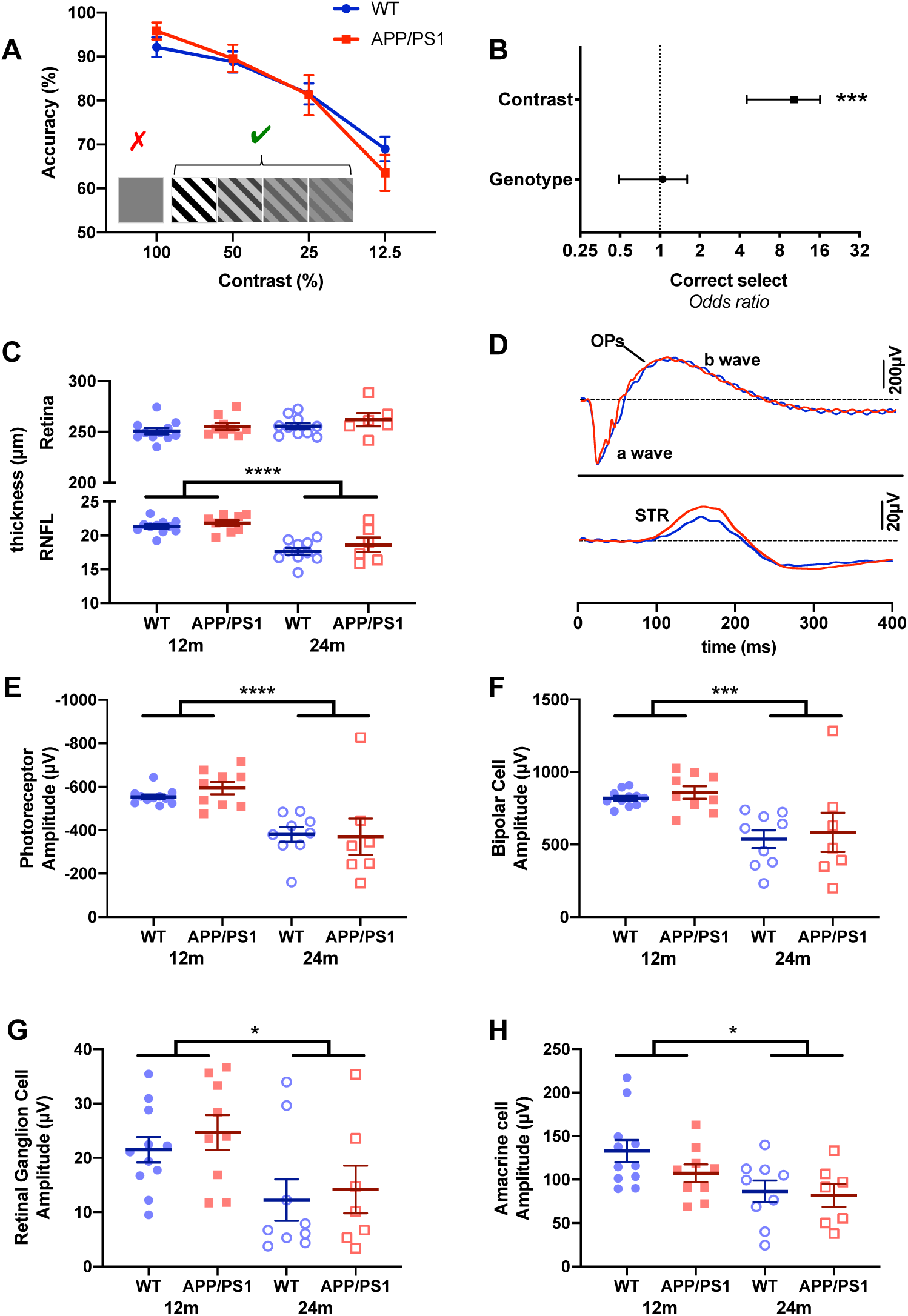
Age but not genotype significantly changes retinal function in 26 month old mice. APP/PS1 mice showed no differences to WT mice on a contrast probe and no degeneration of the retina between 22-26 months. Both APP/PS1 and WT mice were significantly less accurate at lower contrast image identification, but there was no effect of genotype (A, B). There were no changes in total retinal thickness (C) in 12 or 26-month-old WT or APP/PS1 mice (C), but the retinal nerve fibre layer was significantly thinner in 26-month-old mice compared with 12-month-old mice, irrespective of genotype (C). WT and APP/PS1 mice also underwent electroretinography, with representative traces shown in (D). The a-wave represents photoreceptor activity, the b-wave bipolar cell activity, the STR retinal ganglion cell activity and the OP amacrine cell activity. Both 26-month-old WT and APP/PS1 mice showed a reduced amplitude of responsivity of photoreceptors (E), bipolar cells (F), retinal ganglion cells (G) or cells in the inner retina (H) compared to 12-month-old animals, with each dot representing the average of the two eyes per animal. A, C and E-H show mean ± SEM while B shows odds ratios ± 95% CI. TS= touchscreen. *p<0.05, p<0.01, *** p<0.001,**** = p < 0.0001

Both these assessments were applied to 26-month-old mice that had been subjected to touchscreen testing and also a separate naïve cohort to allow the effect of touchscreen training on vision to be assessed. OCT images were analysed for retinal nerve fibre layer (RNFL) and total retinal thickness, with thinner measurements indicating cell loss. 26-month-old mice showed equivalent retinal thickness to 12-month-old mice (Figure 4C, F(2,31) = 2.39, p = 0.13) but significantly thinner RNFL (Figure 4C, F(2,31) = 40.84, p < 0.0001). There was no effect of genotype on total retinal or RNFL thickness (Figure 4C, F(2,31) = 1.34, p = 0.25). When comparing touchscreen trained and behaviourally naïve animals, no significant differences were seen due to either genotype (Supplementary Figure 4A, F(2,25) = 0.22, p = 0.81) or previous touchscreen exposure (Supplementary Figure 4A, F(2,25) = 0.31, p = 0.74), suggesting that both genotype and touchscreen exposure do not cause thinning of the RNFL or the total retina.

Electroretinography (ERG) involved exposing the eyes of mice to short flashes of light and measuring the change in field potential generated in the retina. The health of various cells/layers can be ascertained by analysing parts of the resulting trace (representative traces both WT and APP/PS1 animals is presented in Figure 4D). The a-wave can be taken as a measure of photoreceptor activity, the b-wave as bipolar cell activity, the scotopic threshold response (STR) as retinal ganglion cell activity and finally the oscillatory potential (OP) as a measure of inner retinal activity involving amacrine cell inhibitory neurons. 12-month-old mice showed larger responses than 26-month-old mice in photoreceptor (F(4,27) = 21.35, p < 0.0001), bipolar cell (F(4,27) = 14.17, p = 0.001), retinal ganglion cell (F(4,27) = 6.01, p = 0.020) and amacrine cell responses (F(4,27) = 5.95, p = 0.021). Thus, as animals age, retinal function declines. This decline appears to be genotype independent, with no effect of genotype on amplitude of photoreceptor (F(4,27) = 0.001, p = 0.98), bipolar cell (F(4,27)=0.134, p=0.716), retinal ganglion cell (F(4,27) = 0.75, p = 0.39) or amacrine cell responses (F(4,27) = 2.31, p = 0.14). Similarly, when comparing behaviourally naïve WT and APP/PS1 mice, the magnitude of response of retinal cells was unchanged by genotype (Supplementary Figure 4C-F, photoreceptors: F(4,23) = 0.22, p = 0.65, bipolar cells: F(4,23) = 0.001, p = 0.98, retinal ganglion cells: F(4,23) = 0.69, p = 0.41, amacrine cells: F(4,23) = 0.03, p = 0.86) or touchscreen exposure (Supplementary Figure 4C-F, photoreceptors: F(4,23) = 0.36, p = 0.55, bipolar cells: F(4,23)=1.09, p=0.31, retinal ganglion cells: F(4,23) = 3.62, p = 0.068, amacrine cells: F(4,23) = 3.36, p = 0.78). Thus, contrary to previous reports, we find no evidence of impaired vision in 20-26 month old APP/PS1 mice, indicating the observed behavioural changes in the APP/PS1 mice are driven by cognitive, rather than visual, dysfunction. Furthermore, prolonged touchscreen exposure does not change retinal health as measured by OCT or ERG.

## Discussion

In this study, we have shown a progressive executive function deficit in APP/PS1 mice, with intact performance in the reversal learning task at 13 months of age and a deficit appearing by 22 months of age. This impairment was independent of changes in pairwise discrimination and any genotype-mediated structural and functional changes to the retina. Extinction learning was assessed at 13, 16 and 26 months of age, and was found to first be impaired by 16 months. This is the first demonstration of an appetitive executive function deficit in the APP/PS1 mouse and further validates the use of touchscreens as a method for assessing cognition in pre-clinical AD models.

APP/PS1 animals have previously shown deficits in reversal learning. In reversal MWM maze tasks, APP/PS1 mice have shown acquisition deficits from 8 (but not 6) months relatively consistently in both male and female mice (Gallagher, Minogue, & Lynch, 2012; Hooijmans et al., 2009; Jankowsky et al., 2005; Savonenko et al., 2005; Stover & Brown, 2012), although one study identified acquisition deficits in female mice only (Gallagher et al., 2012). Furthermore, 6-9 month old APP/PS1 mice have shown deficits in a left-right discrimination reversal task in the T water maze (Filali et al., 2009) and this deficit was rescued with memantine (Filali et al., 2011). However, these studies utilise cold-water stress to motivate animal behaviour, and stress has been shown to negatively affect cognition (reviewed in (Joëls & Baram, 2009), thus it is important to consider rewarded reversal tasks in addition to aversively motivated ones. To our knowledge, there is only one study that evaluates reversal in a cheeseboard task motivated by reward, which showed that 6-8 month old APP/PS1 male mice were not impaired on acquisition of the reversal task (Cheng et al., 2013). This, in addition to our own finding that APP/PS1 mice were unimpaired at 13 months but impaired by 22 months in rewarded touchscreen reversal, indicates that the age at which APP/PS1 mice show reversal deficits in aversive compared to rewarded reversal tasks may be different.

There is precedent for impairments in appetitive reversal in AD mouse models other than our study. Tg2576 animals, which have the same Swedish mutation in APP as the APP/PS1 mice are driven by a similar PrP promoter (but lack the PS1 transgene), have shown deficits in rewarded reversal digging tasks. In these tasks, mice are required to learn to dig in one of two essential-oil-scented pots for a food reward. Tg2576 mice have shown impaired reversal at 6 (Zhuo et al., 2008, 2007) and 12 months (Shirey et al., 2009) on these tasks. Interestingly, the digging reversal deficit seen at 6 months in Tg2576 mice was seen without increased perseveration (Zhuo et al., 2008), which contrasts with our study, where the 22-25 month old APP/PS1 animals showed a significant increase in perseveration. Another interesting aspect of one of these studies is that the reversal deficit in Tg2576 mice observed at 6 months was not apparent at 14 months - this was due to 14-month-old WT control animals showing reduced reversal performance, to the same level of performance observed in 6 and 14-month-old Tg2576 mice (Zhuo et al., 2007). This indicated that perhaps aging out to 14 months causes age-dependent cognitive decline in WT animals, and that this cognitive decline could mask reversal deficits in AD models that are apparent earlier in life. We observed an aging effect in our 22 month WT animals compared to 13 month WT animals in both PD and Reversal task, but at 22 months the decreased performance in PD and reversal attributable to aging is significantly smaller than the AD deficit. Future studies could focus on disentangling aging and AD effects.

In considering deficits in touchscreen reversal, three AD mouse models have been run on the same PD and reversal touchscreen task used in this study. 4-5 month old APPSwDI/*Nos2-/-* mice were run on 20 days of the reversal and showed an attenuated acquisition of reversal, as well as increased errors, analogous to the deficits seen in our 22-25 month old APP/PS1 mice (Piiponniemi et al., 2017). However, as the APPSwDI/*Nos2-/-* animals were not run to criterion, it is unclear if they would ever be able to acquire the task (Piiponniemi et al., 2017). Additionally, the APPSwDI/*Nos2-/-* model has an artificially accelerated phenotype due to the homozygous null mutation of the Nos2 gene (which has anti-apoptotic and pro-survival functions). Another model, the APPPS1-21 mouse model, which has the Swedish mutation in the APP gene and the L166P point mutation in the PS1 gene, was shown to have unimpaired PD but reduced accuracy in reversal compared to their WT counterparts at 6 and 9 months. However, the APPPS1-21 animals performed significantly less trials throughout reversal, thus it is not possible to ascertain if this is a learning or motivational deficits. Conversely, TgCRND8 mice showed accelerated, rather than attenuated, acquisition of reversal in the touchscreen paradigm. This deficit was not due to weaker stimulus-reward association as WT and TgCRND8 showed equal retention memory of the rewarded stimulus 10 days after finishing reversal (Romberg et al., 2013). This acceleration is counter to what would be expected from the clinical literature. In patients, cognitive flexibility is assessed by card-sorting tasks, where patients are required to inhibit a response to an initially learned rule and learn a new rule - similar to what is required of animals in reversal. AD patients show attenuated acquisition of set-shifting rules and increased perseveration errors on card-sorting tasks (Andriuta et al., 2018; Baudic et al., 2006; Huang et al., 2017; Ramanan et al., 2017). Thus, the human data mirrors our observed attenuated reversal acquisition and increased perseverative behaviour in APP/PS1 mice at 21-23 months of age. This is the first time a reversal deficit mirroring those seen in humans has been shown to develop following a period of non-impairment in an aged, AD mouse model.

Extinction deficits have also been observed in APP/PS1 mice in aversive extinction tasks (Bonardi et al., 2011; Cheng et al., 2019; Ramírez-Lugo et al., 2009). However, like reversal, deficits in rewarded and aversive extinction appear to dissociate in APP/PS1 animals. Bonardi et al. (2011) demonstrated attenuation of contextual fear conditioned extinction but unaffected auditory-driven appetitive extinction in 4-month-old female APP/PS1 mice. In our study, we saw an attenuation of extinction in male APP/PS1 mice at 16 but not 13 months of age. There are multiple reasons why we see a deficit in our animals compared to previous studies (Bonardi et al., 2011). First, and most significantly, our animals are notably older, and as this is a progressive model, the animals are much more likely to show a deficit at a later age. Secondly, the procedure used was slightly different, with our procedure requiring animals to complete 30 trials in every acquisition and extinction session, while Bonardi et al. (2011) used a 15 trial acquisition and 10 trial extinction. As the number of trials is increased in our paradigm, and all animals performed multiple touchscreen tasks prior to extinction, it is likely the animals develop a stronger stimulus/reward association. Furthermore, the fact animals are exposed to more trials in the touchscreen paradigm allows us to pick up more subtle changes. Regardless of the reason, this is the first study to show deficits in rewarded extinction in APP/PS1 mice.

This is not the first study to show changes in touchscreen extinction in an AD mouse model. Deficits in touchscreen extinction have been shown previously in the TgCRND8 mice. As with reversal, TgCRND8 mice show accelerated extinction, in contrast with the attenuation seen here in the APP/PS1 mice. Extinction is not actively tested in humans, but like reversal, it relies on the PFC function, and as such we would anticipate similar patterns in extinction as reversal or set shifting behaviour. A possible explanation for the accelerated extinction and reversal seen in the TgCRND8 mice compared to the attenuation seen in APP/PS1 mice (and more broadly, human patients) lies in sub regions of the PFC. Interestingly, lesions in the medial PFC have been shown to cause accelerated extinction and reversal (Graybeal et al., 2011), while lesions in the orbitofrontal and infralimbic cortex, two other parts of the PFC, impair reversal (Chudasama & Robbins, 2003). Thus, it could be that 4-5 month old TgCRND8 mice have more specific impairments in the medial PFC, while the 22-25 month old APP/PS1 mice have wider PFC dysfunction which accounts for the slower acquisition of extinction and reversal in the APP/PS1 animals (Izquierdo & Murray, 2005; Malá et al., 2015). As such, the opposite pattern in these studies is still consistent with wider PFC dysfunction seen in AD.

However, once a deficit manifests one would not expect it to resolve – so why don’t 26-month-old APP/PS1 mice show an attenuation of extinction? Both 26-month-old WT and APP/PS1 mice appear to have struggled with the time restrictions in the extinction. First, 26-month-old animals took longer to reach criterion on the acquisition stage, with 2 animals never reaching criterion at all, as opposed to 13 or 16-month-old animals. Furthermore, response latencies were longer in 26-month animals and, finally, response rates on day 1 of extinction were about 60% compared to 80-90% in 13 and 16-month-old animals. This low response rate may have induced a floor effect, masking any potential deficits.

What could account for the intact performance of 12-13 month old APP/PS1 mice, apart from the already discussed dissociation of rewarded and aversive EF tasks? First, it may be that these animals are simply not old enough. Previously, a systematic review comparing 49 behavioural studies in the Tg2576 mouse model indicated the age of onset of cognitive deficits was relatively variable (Stewart, Cacucci, & Lever, 2011). Secondly, another factor is that all animals in the present study were food restricted in order to motivate touchscreen task performance. Long-term caloric restriction (CR) is known to delay onset of MWM acquisition deficits (Halagappa et al., 2007) and reduced amyloid beta levels in AD mouse models (Halagappa et al., 2007; Mouton, Chachich, Quigley, Spangler, & Ingram, 2009; Patel et al., 2005; Schafer et al., 2015; Wang et al., 2005) and normal aging squirrel monkeys (Qin et al., 2006). Finally, CR also improved cognitive outcomes in obese patients with MCI (Horie et al., 2016). A study looking at the effect of a 4 month 40% food restriction on male 13-14 month APP/PS1 mice, showed a 30% reduction in Aβ levels in the hippocampus and neocortex (Mouton et al., 2009). In our study, the 13-month cohort of animals started food restriction and touchscreen testing at 6-7 months while our 16 month cohort started at 8-9 months. Thus, while the 40% CR is much stricter than the 15-25% used in this study, it is likely that caloric restriction reduced amyloid in our animals, and potentially improved cognition as seen in 3xTG-AD mice in another CR study (Halagappa et al., 2007). Again, future studies are required to investigate the effect of mild, long-term CR and how this attenuates cognitive deficit development, as this has never been done in an AD mouse model.

The fact that APP/PS1 animals show no deficit in PD up to 22 months is similar to previous studies. Both TgCRND8 (Romberg et al., 2013) and APPSwDI/*Nos2-/-* (Piiponniemi et al., 2017) mice showed no changes in acquisition of PD on the same touchscreen task. However, other studies have shown PD deficits in APP/PS1 mice in discrimination paradigms at a younger age than this. Attenuated discrimination learning in a home cage in a 3-hole choice paradigm was shown in 6-7 month old male APP/PS1 mice (Remmelink, Smit, Verhage, & Loos, 2016), indicating that even basic learning can be affected in this strain. However, these animals are on a pure C57BL/6J background, and genetic diversity in AD mouse models have been shown to drastically change phenotype (Neuner, Heuer, Huentelman, O’Connell, & Kaczorowski, 2019), thus potentially explaining this difference. Furthermore, the touchscreen paradigm is evidently different to a home-cage learning paradigm. Touchscreen testing requires the animals to be handled, and food restricted, and this causes many changes in the brains of touchscreen-tested animals (Mallien et al., 2016). These changes could also potentially explain this deficit dissociation between basic learning in the home cage and the touchscreens. Thus, intact PD in touchscreens seems consistent across multiple AD mouse models, and allows us to be confident that the reversal deficit is specific to cognitive flexibility rather than any issues in basic learning.

We assessed APP/PS1 animals late in disease progression for visual deficits in a touchscreen contrast task, and also scrutinised visual structure and function using optical coherence tomography (OCT) and electroretinography (ERG). Using these approaches, we did not see any difference in vision between APP/PS1 and WT mice, although clear aging effects were seen. The age-related changes in retinal function and structure seen here are consistent with those previously documented for both mice (Kolesnikov, Fan, Crouch, & Kefalov, 2010; Shariati, Park, & Liao, 2015; Vessey et al., 2015) and rats (Nadal-Nicolás, Vidal-Sanz, & Agudo-Barriuso, 2018). The current study highlights that at a very advanced age there is thinning of the retinal nerve fibre layer, without a change in total retina thickness (Figure 4C). Deficits in all ERG components in 26-month-old WT mice (Figure 4E-4H) suggest that there is impaired neuronal function, that is greater than expected from age-related neuronal loss (Nadal-Nicolás et al., 2018), as indicated by the maintenance of total retinal thickness (Figure 4C).

Amyloid beta has been shown to build up in the retina of male and female APP/PS1 animals (Gupta et al., 2016; Koronyo-Hamaoui et al., 2011; Ning, Cui, To, Ashe, & Matsubara, 2008; Perez, Lumayag, Kovacs, Mufson, & Xu, 2009) and changes in a-wave and b-wave activity following ERG at 12-16 months (Perez et al., 2009) and scotopic threshold response at 13-16 months (Gupta et al., 2016) have been shown in the same mouse model with the same genetic background, neither of which we replicated here. Interestingly a study which assessed 3 – 12 month old APP/PS1 mice found a lack of retinal pathology as assessed with immunohistochemistry, western blot and qPCR techniques (Chidlow, Wood, Manavis, Finnie, & Casson, 2016). Alterations to thicknesses in retinal ganglion cell OCT measures such as retinal nerve fibre layer are suggested to be more sensitive than other retinal layers to AD related changes in both human (Armstrong, 2009; Lim et al., 2016) and animal models (Liu et al., 2009); despite this, we saw no changes in RNFL thickness in our APP/PS1 mice. Furthermore, (Stover & Brown, 2012) tested the same background strain of pooled male and female APP/PS1 animals at 5-8 or 20-26 months in the visible water box on a series of visual discrimination and acuity tests. When animals had to distinguish between horizontal and vertical lines to find the escape platform, old mice showed virtually no improvements over 8 days, regardless of genotype, unlike here, where all 21-22 month old WT mice acquired PD. Furthermore, when Stover and Brown (2012) assessed visual acuity by increasing the cycles per degree of the vertical lines (i.e. increasing the number of lines in the same spaces) compared to a grey square, the old WT mice performed better than old APP/PS1 animals indicating that the APP/PS1 mice had poorer vision; in our study, both APP/PS1 and WT mice performed equally on the contrast probe task. However, in these old animals, female mice perform worse than male mice regardless of genotype (Stover & Brown, 2012), and in this study we used only male mice. It is possible that the effects mentioned above are driven by differences in female mice, with male mice showing little change, thus accounting for lack of changes in ERG and OCT measurements between genotypes in this study. However, AD patients report visual deficits, with many measurable changes in the eye appearing through the disease (reviewed in Armstrong 2009; Lim et al. 2016). Specifically, RNFL thinning has been shown in living AD patients by OCT (Berisha, Feke, Trempe, McMeel, & Schepens, 2007; La Morgia et al., 2016; Liu et al., 2015) and marked axonal and retinal ganglion cell loss in post-mortem tissue (La Morgia et al., 2016). Large concentrations of Aβ in and around retinal ganglion cells have also been noted (La Morgia et al., 2016). Of note, increased Aβ is also found in visual areas of the brain both in normal aging of humans (Sepulcre et al., 2017) and animals (Hernández-Zimbrón et al., 2017), as well as AD patients (Rowe et al., 2013) and AD mouse models like the APP/PS1 (Whitesell et al., 2018). Additionally, dysfunction of the visual areas has been shown in mild AD patients (Brewer & Barton, 2014), thus visual changes may not be driven by the eye alone. However, we found no evidence of behavioural changes due to visual impairment in the contrast touchscreen task. This, together with the data from the ERG and OCT, indicates that vision is intact in APP/PS1 mice used in this study. It is unclear why this differs from previous studies in APP/PS1 mice that showed RNFL thinning and changes in a-, b- and STR wave activity, and more research needs to be conducted to understand why these changes were not recapitulated in this study.

In summary, we have shown reversal learning impairment in 22-25 month old but not 12-13 month old APP/PS1 mice using a reward-based touchscreen task. We have also shown impairments in extinction by 16 months of age, supporting the idea of broader executive dysfunction in this mouse model. We have shown that these impairments occur without any changes in vision at the same time-points. The reversal impairment is in line with many other reports using exploratory and aversive motivated reversal tasks with the APP/PS1 mouse model; however the impairment described in this study occurs at a much later age, indicating that the touchscreen paradigm may not be suited for early detection of executive dysfunction. Extinction impairments identified in this study also occur later than previous reports. An alternative explanation is that the lengthy touchscreen testing that occurs in order for animals to acquire the task could act in a similar way as computerised cognitive training does in humans. Furthermore, mice are calorie restricted in order to motivate them to perform in touchscreens. One CR study has shown rescued performance in the MWM in the 3xTG-AD mouse model of AD, but no other studies have evaluated the effect of CR on cognition in AD mouse models. More research is needed in order to understand the effect of long-term touchscreen training and mild CR on cognition in AD mouse models. The present study adds to the growing literature supporting the utility of touchscreens to identify and quantify executive function deficits in a clinically translatable way and provides a method to screen cognitively enhancing treatments targeted at executive function. Considering executive function seems to predict cognitive decline and MCI to AD conversion (Mez et al., 2013), this is a particularly important domain to measure both pre-clinically and track clinically in AD.

## Acknowledgements

We would like to thank past and present laboratory members for useful discussions and technical assistance. We thank Britany Cuic, Maddison Ible, Daniel Drieberg, Shannon Currin Craig Thompson and Brett Purcell for their assistance in mouse husbandry and management of equipment. Thank you to Nippy’s Ltd, for the donation of Ice Strawberry milk to our study. A.S is supported by an Australian Government Research Training Program Scholarship and Yulgilbar top-up scholarship. A.M.Z.J is supported by an Australian Government Research Training Program Scholarship A.J.H. is a National Health and Medical Research Council (NHMRC) Principal Research Fellow and has been supported by an Australian Research Council (ARC) FT3 Future Fellowship (FT100100835). E.L.B. is supported by a NHMRC-ARC Dementia Research Development Fellowship. C.T.O.N is supported by an Australian Research Council (ARC) Linkage grant (LP160100126) and a Melbourne Research Fellowship. J.K.H.L. is supported by the Guelma-Alaexander fellowship in Neuroscience and V.H.Y.W. are supported by an Australian Research Council (ARC) Linkage grant (LP160100126). This work was supported by directed research costs associated to a NHMRC-ARC Dementia Research Development fellowship. We would also like to acknowledge the operational infrastructure support from the State Government of Victoria.

