## Supplementary Material for "Progressive impairments in executive function in the APP/PS1 model of Alzheimer’s disease as measured by translatable touchscreen testing"

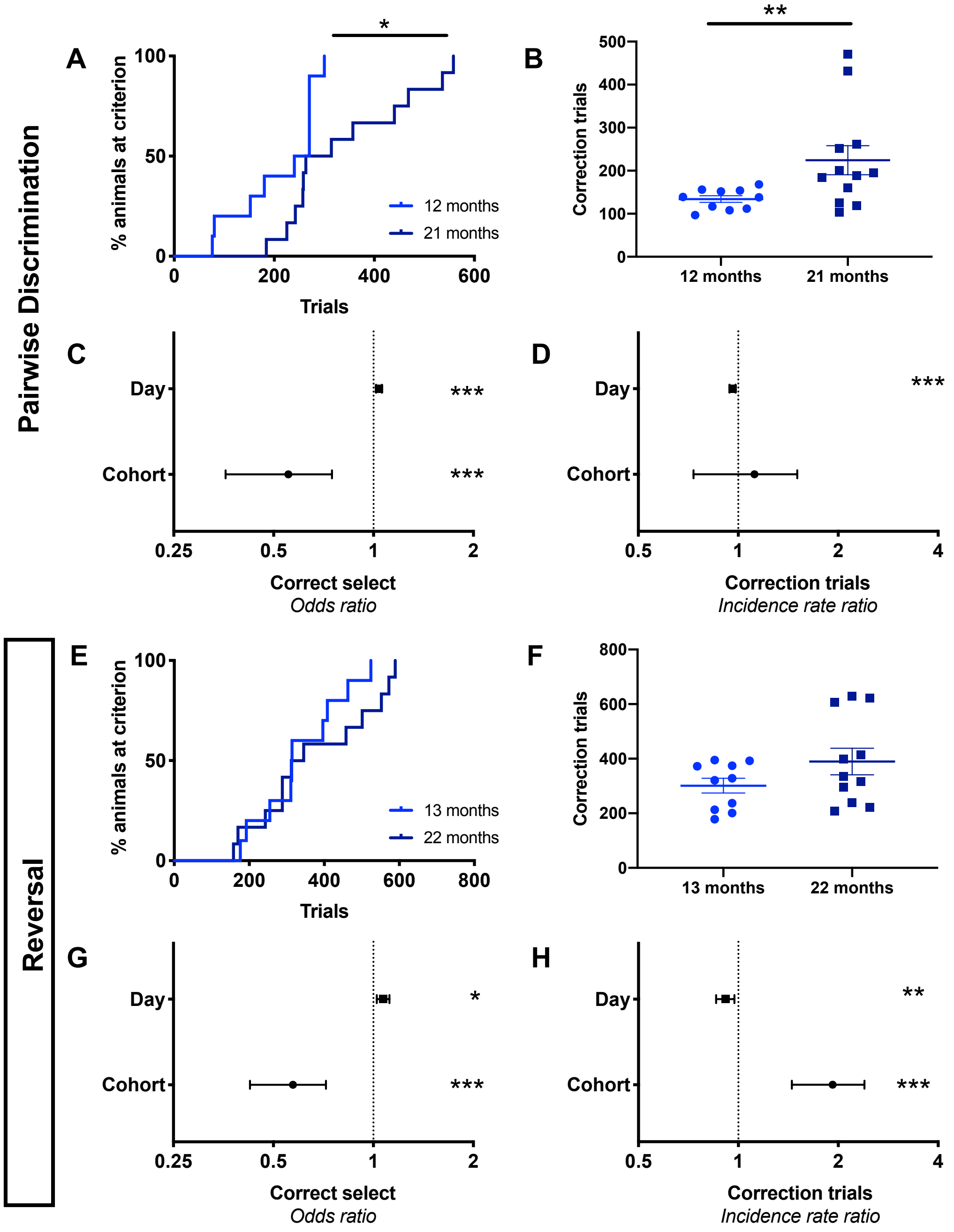


**Supplementary Figure 1. 21-22 month old WT mice showed attenuated PD and Reversal compared to 12-13 month old mice.** 21-22 month old mice were impaired across multiple measures on PD and reversal compared to 12-13 month old mice. 21-month-old WT mice took an increased number of trials to reach criterion for PD (A), and were less likely to choose the correct image given any unique trial (C) compared to 12-month-old WT mice. Additionally, 21-month-old WT mice took more correction trials to reach criterion (B) but showed the same incidence rate ratio of correction trials (D) as 13-month-old WT mice in PD. In reversal, 13- and 22- month old WT mice completed an equivalent number of trials to criterion (E) and correction trials (F). 22-month-old WT mice show a decreased odds ratio of correct selection (G) and an increased expected correction trial count per incorrect trial as 13-month-old WT animals during reversal (H). A and C show survival curves, B and F show mean ± SEM while C-D and G-H show odds ratios or incidence rate ratios ± 95% CI. * = $p < 0.05$; ** = $p < 0.01$; *** = $p < 0.001$

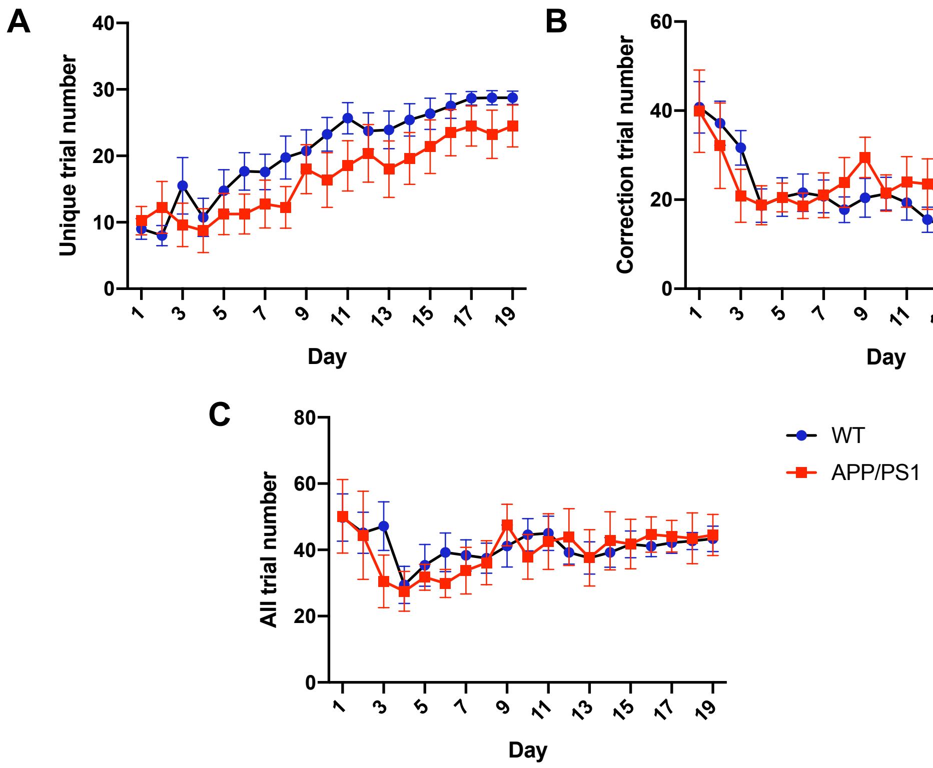


**Supplementary Figure 2. 22-month-old APP/PS1 and WT mice complete the same number of trials during reversal.** 22-month-old APP/PS1 mice completed a similar number of trials in the first 19 days of reversal compared to WT mice. 22-month-old APP/PS1 mice completed a similar number of unique trials (A) correction trials (B) and combined trials (C) as WT animals. A-C show group means ± SEM, * = $p < 0.05$; *** = $p < 0.001$

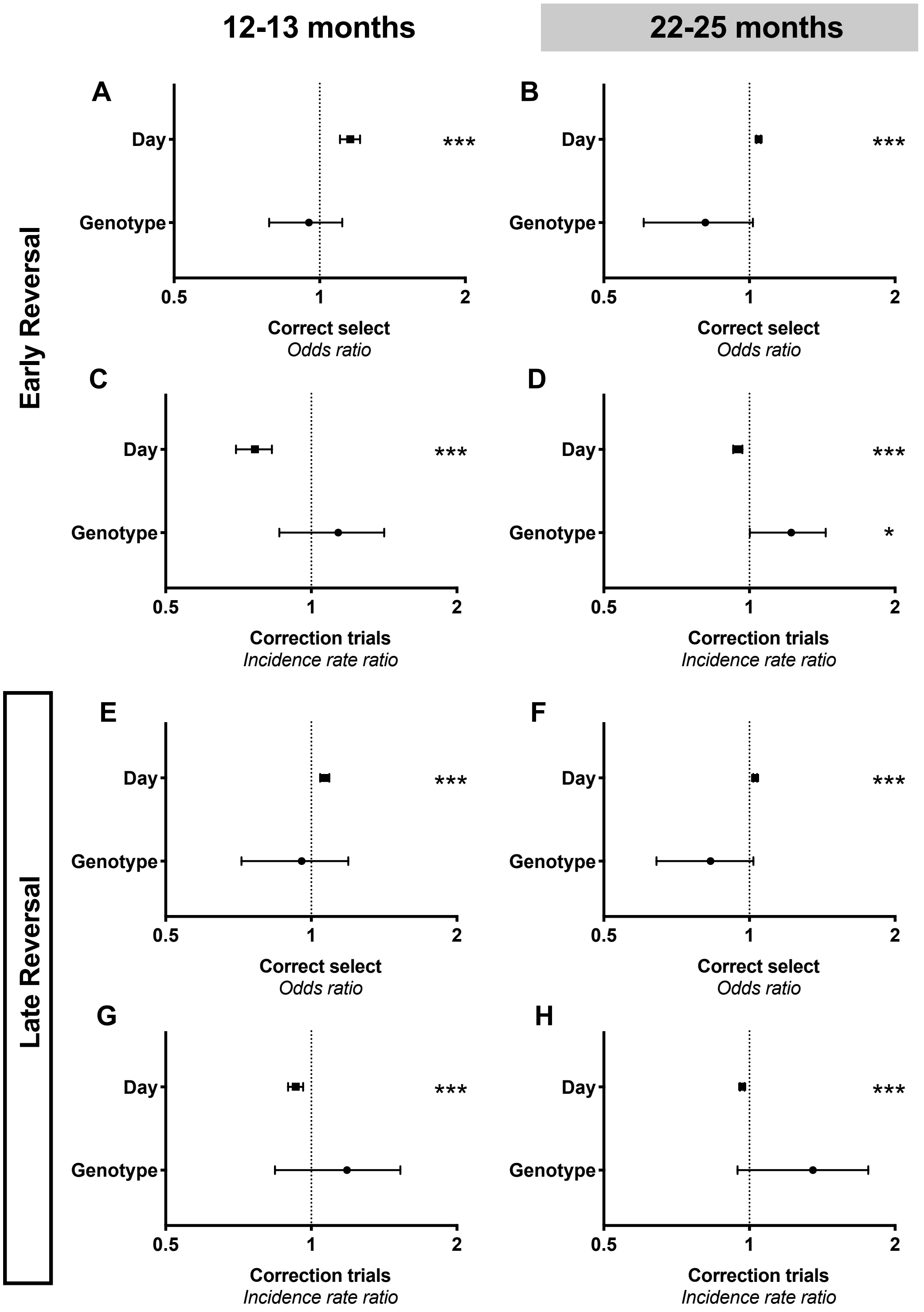


**Supplementary Figure 3. Early and late reversal was attenuated in 22-month-old APP/PS1 mice but unaffected in 13-month-old APP/PS1 animals.** 22-month-old mice showed similar trends to overall reversal in early and late reversal learning while 13-month-old mice were unaffected on all measures. 13-month-old APP/PS1 mice were just as likely to choose the correct image given any unique trial in both early (A) and late (E) reversal. 22-month-old APP/PS1 mice are non-significantly less likely to choose the correct image given any unique trial during early (B) and late (D) reversal. 13-month-old APP/PS1 mice showed the same expected correction trial count per incorrect trial as WT animals during early (C) and late (G) reversal. 22-month-old APP/PS1 mice performed more correction trials per incorrect trial than WT animals during early reversal (D), with a similar but non-significant trend during late reversal (H). A-B and E-F show odds ratios ± 95% CI while C-D and G-H show incidence risk ratios ± 95% CI. * = $p < 0.05$; *** = $p < 0.001$

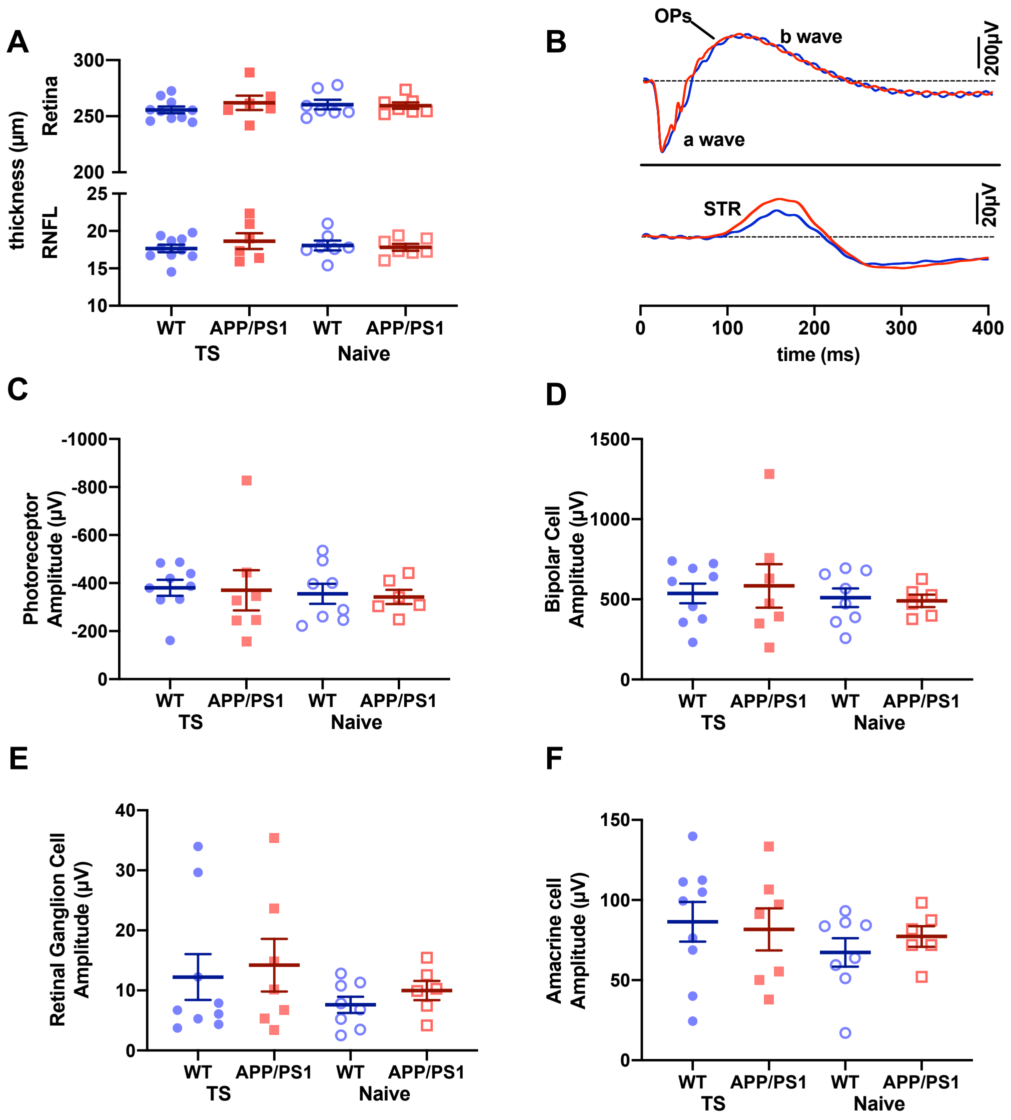


**Supplementary Figure 4. Touchscreen training has no effect on retinal structure and function.** Behaviourally naïve and touchscreen tested APP/PS1 and WT mice shown similar OCT and ERG measures. 20-26 month old behaviourally naive mice show equivalent RNFL and total retinal thickness as 26-month-old touchscreen trained WT and APP/PS1 mice as measured by OCT (A). A typical ERG trace shows similar waveforms in APP/PS1 and WT mice (B) Behaviourally naive and touchscreen trained APP/PS1 and WT mice show similar magnitude amplitude changes in photoreceptors (C), bipolar cells (D), retinal ganglion cells (E) and amacrine cells (F). A and C-F show mean ± SEM, * = $p < 0.05$; *** = $p < 0.001$
